# The HTM Spatial Pooler – a neocortical algorithm for online sparse distributed coding

**DOI:** 10.1101/085035

**Authors:** Yuwei Cui, Subutai Ahmad, Jeff Hawkins

**Affiliations:** Numenta, Inc, Redwood City, California, USA

## Abstract

Hierarchical temporal memory (HTM) provides a theoretical framework that models several key computational principles of the neocortex. In this paper we analyze an important component of HTM, the HTM spatial pooler (SP). The SP models how neurons learn feedforward connections and form efficient representations of the input. It converts arbitrary binary input patterns into sparse distributed representations (SDRs) using a combination of competitive Hebbian learning rules and homeostatic excitability control. We describe a number of key properties of the spatial pooler, including fast adaptation to changing input statistics, improved noise robustness through learning, efficient use of cells and robustness to cell death. In order to quantify these properties we develop a set of metrics that can be directly computed from the spatial pooler outputs. We show how the properties are met using these metrics and targeted artificial simulations. We then demonstrate the value of the spatial pooler in a complete end-to-end real-world HTM system. We discuss the relationship with neuroscience and previous studies of sparse coding. The HTM spatial pooler represents a neurally inspired algorithm for learning sparse representations from noisy data streams in an online fashion.

## 2. Introduction

Our brain continuously receives vast amounts of information about the external world through peripheral sensors that transform changes in light luminance, sound pressure, and skin deformations into millions of spike trains. Each cortical neuron has to make sense of a flood of time-varying inputs by forming synaptic connections to a subset of the presynaptic neurons. The collective activation pattern of populations of neurons contributes to our perception and behavior. A central problem in neuroscience is to understand how individual cortical neurons learn to respond to specific input spike patterns, and how a population of neurons collectively represents features of the inputs in a flexible, dynamic, yet robust way.

Hierarchical temporal memory (HTM) is a theoretical framework that models a number of structural and algorithmic properties of the neocortex (Hawkins et al., 2011). HTM networks can learn time-based sequences in a continuous online fashion using realistic neuron models that incorporate nonlinear active dendrites (Antic et al., 2010; Major et al., 2013) with thousands of synapses (Hawkins and Ahmad, 2016). When applied to streaming data, HTM networks achieve state of the art performance on anomaly detection (Lavin and Ahmad, 2015; Ahmad and Purdy, 2016) and sequence prediction tasks (Cui et al., 2016a).

The success of HTM relies on the use of sparse distributed representations (SDRs) (Ahmad and Hawkins, 2016). Such sparse codes represent a favorable compromise between local codes and dense codes (Földiák, 2002). It allows simultaneous representation of distinct items with little interference, while still has a large representational capacity (Kanerva, 1988; Ahmad and Hawkins, 2015). Existence of SDRs have been documented in auditory, visual and somatosensory cortical areas (Vinje and Gallant, 2000; Weliky et al., 2003; Hromádka et al., 2008; Crochet et al., 2011). HTM spatial pooler (SP) is a key component of HTM networks that continuously encodes streams of sensory inputs into SDRs. Originally described in (Hawkins et al., 2011), the term “spatial pooler” is used because input patterns that share a large number of co-active neurons (i.e. that are spatially similar) are grouped together into a common output representation. Recently there has been increasing interest in the mathematical properties of the HTM spatial pooler (Pietroń et al., 2016; Mnatzaganian et al., 2017) and machine learning applications based on it (Thornton and Srbic, 2011; Ibrayev et al., 2016). In this paper we explore several functional properties of the HTM spatial pooler that have not yet been systematically analyzed.

The HTM spatial pooler incorporates several computational principles of the cortex. It relies on competitive Hebbian learning (Hebb, 1949), homeostatic excitability control (Davis, 2006), topology of connections in sensory cortices (Udin and Fawcett, 1988; Kaas, 1997) and activity-dependent structural plasticity (Zito and Svoboda, 2002). The HTM spatial pooler is designed to achieve a set of computational properties that support further downstream computations with SDRs. These properties include (1) preserving the semantic similarity of the input space by mapping similar inputs to similar outputs, (2) continuously adapting to changing statistics of the input stream, (3) forming fixed sparsity representations, (4) being robust to noise, and (5) being fault tolerant. As an integral component of HTM, the outputs of the SP can be easily recognized by downstream neurons and contribute to improved performance in an end-to-end HTM system.

The primary goal of this paper is to provide a thorough discussion of the computational properties of the HTM spatial pooler and demonstrate its value in end-to-end HTM systems. The paper is organized as follows. We first introduce the HTM spatial pooler algorithm. We discuss the computational properties in detail and then describe a set of metrics to quantify them. We demonstrate how these properties are satisfied, first using specific isolated simulations, and then in the context of an end-to-end HTM system. In the discussion, we propose potential neural mechanisms and discuss the relationship to existing sparse coding techniques.

## 3. Model

The spatial pooler is a core component of HTM networks (Fig. 1A). In an end-to-end HTM system, the spatial pooler transforms input patterns into SDRs in a continuous online fashion. The HTM temporal memory learns temporal sequences of these SDRs and makes predictions for future inputs (Cui et al., 2016a; Hawkins and Ahmad, 2016). A single layer in an HTM network is structured as a set of mini-columns, each with a set of cells (Fig. 1B). The HTM neuron model incorporates dendritic properties of pyramidal cells in neocortex (Spruston, 2008), where proximal and distal dendritic segments on HTM neurons have different functions (Fig. 1C) (Hawkins and Ahmad, 2016). Patterns detected on proximal dendrites lead to action potentials and define the classic receptive field of the neuron. Patterns recognized by a neuron’s distal synapses act as predictions by depolarizing the cell without directly causing an action potential.

**Figure 1.**
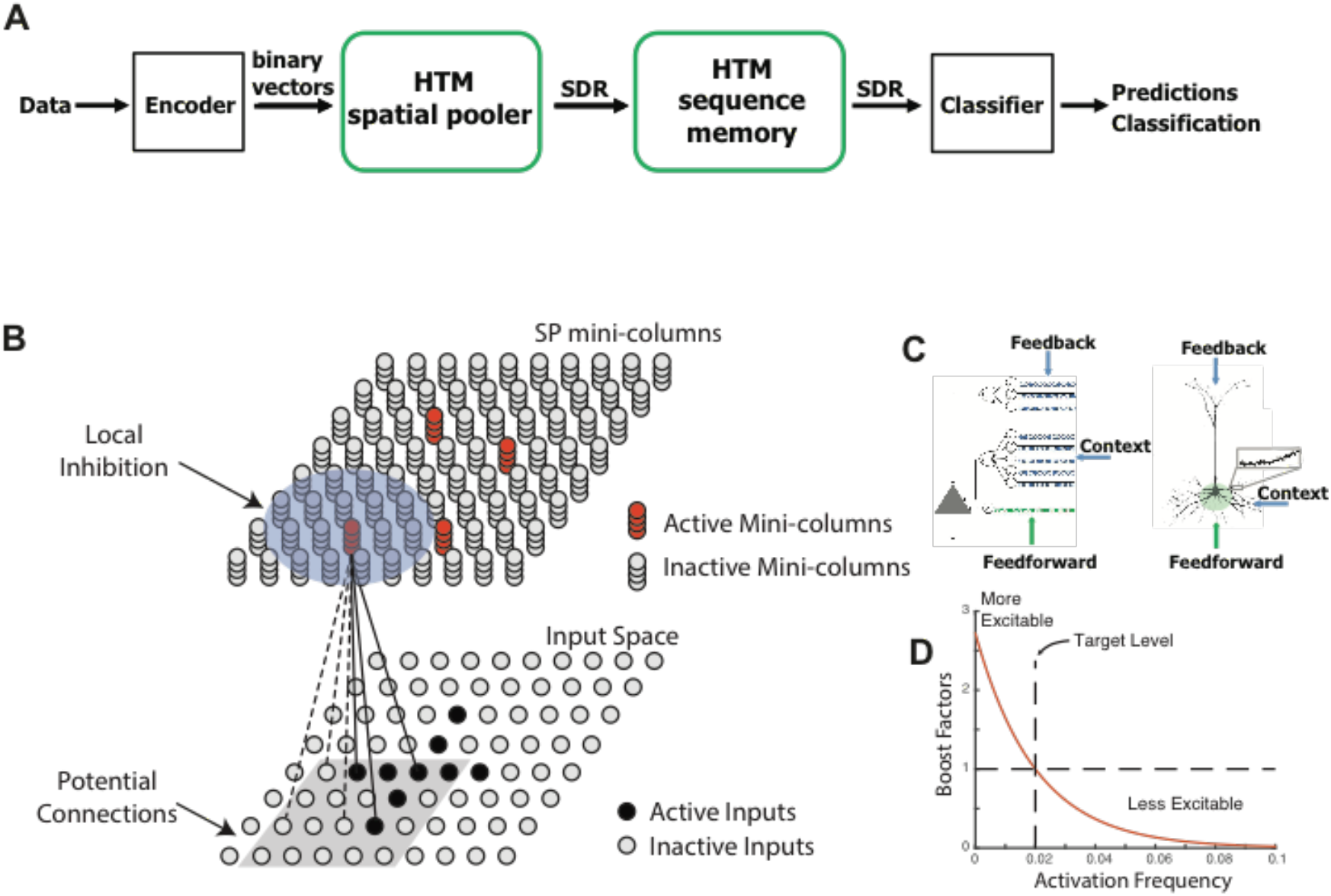
HTM Spatial Pooler. **A.** An end-to-end HTM system consists of an encoder, the HTM spatial pooler, the HTM temporal memory and an SDR classifier. **B.** The HTM spatial pooler converts inputs (*bottom*) to SDRs (*top*). Each SP mini-column forms synaptic connections to a subset of the input space (gray square, potential connections). A local inhibition mechanism ensures that a small fraction of the SP mini-columns that receive most of the inputs are active within the local inhibition radius (shaded blue circle). Synaptic permanences are adjusted according to the Hebbian rule: for each active SP mini-column, active inputs (black lines) are reinforced and inactive inputs (dashed lines) are punished. **C.** An HTM neuron (left) has three distinct dendritic integration zones, corresponding to different parts of the dendritic tree of pyramidal neurons (right). The SP models the feedforward connections onto the proximal dendrite. **D**. The excitability of a SP mini-column depends on its past activation frequency.

In HTM theory different cells within a mini-column represent this feedforward input in different temporal contexts. The spatial pooler models synaptic growth in the proximal dendritic segments. Since cells in a mini-column share the same feedforward classical receptive field (Buxhoeveden, 2002), the SP models how this common receptive field is learned from the input. The SP output represents the activation of mini-columns in response to feedforward inputs. The HTM temporal memory models a cell’s distal dendritic segments and learns transitions of SDRs by activating different sets of cells depending on the temporal context (Hawkins and Ahmad, 2016). The output of HTM temporal memory represents the activation of individual cells across all mini-columns.

The SP models local inhibition among neighboring mini-columns. This inhibition implements a *k*-winners-take-all computation (Majani et al., 1988; Makhzani and Frey, 2015). At any time, only a small fraction of the mini-columns with the most active inputs become active. Feedforward connections onto active cells are modified according to Hebbian learning rules at each time step. A homeostatic excitatory control mechanism operates on a slower time scale. The mechanism is called “boosting” in (Hawkins et al., 2011), because it increases the relative excitability of mini-columns that are not active enough. Boosting encourages neurons with insufficient connections to become active and participate in representing the input.

Each SP mini-column forms synaptic connections to a population of input neurons. We assume that the input neurons are arranged topologically in an input space. The location of the *j*th input neuron is denoted as **x**_j_. The dimensionality of the input space depends on applications. For example, the input space is 2-dimensional if the inputs are images and 1-dimensional if the inputs are scalar numbers. A variety of encoders are available to deal with different data types (Purdy, 2016). The output neurons are also arranged topologically in a different space; we denote the location of the *i*th SP mini-column as **y**_i_.

We use HTM neuron models in the SP (Fig. 1B). A complete description of the motivation and supporting evidence for this model can be found in (Hawkins and Ahmad, 2016). In this model, the learning rule is inspired by neuroscience studies of activity-dependent synaptogenesis (Zito and Svoboda, 2002). The synapses for the *i*th SP mini-column are located in a hypercube of the input space centered at 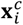 with an edge length of γ. Each SP mini-column has potential connections to a fraction of the inputs in this region. We call these “potential” connections because a synapse is connected only if its synaptic permanence is above the connection threshold. The set of potential input connections for the *i*th mini-column is, 
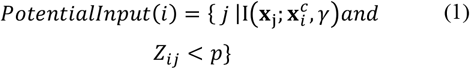

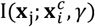 is an indicator function that returns one only if **x**_j_ is located with a hypercube centered at 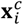 with an edge length of γ.Z_ij_ ~ *U*(0, 1) is a random number uniformly distributed in [0, 1], *p* is the fraction of the inputs within the hypercube that are potential connections.

We model each synapse with a scalar permanence value and consider a synapse connected if its permanence value is above a connection threshold. We denote the set of connected synapses with a binary matrix **W**, 
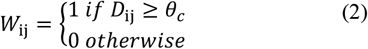

Where *D*_ij_ gives the synaptic permanence from the *j*th input to the *i*th SP mini-column. The synaptic permanences are scalar values between 0 and 1, which are initialized to be independent and identically distributed according to a uniform distribution between 0 and 1 for potential synapses. 
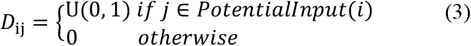

The connection threshold *θ_c_* is set to be 0.5 for all experiments, such that initially 50% of the potential synapses are connected. Performance of the SP is not sensitive to the connection threshold parameter.

Neighboring SP mini-columns inhibit each other via a local inhibition mechanism. We define the neighborhood of the *ith* SP mini-column y_i_ as 
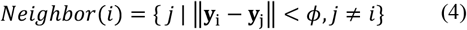

Where ||**y**_i_ − **y**_j_|| is the Euclidean distance between the mini-column *i* and *j*. Since local inhibition occurs among neighboring mini-columns, the parameter *ϕ* controls the inhibition radius. Local inhibition is important when the input space has topology, that is, when neighboring input neurons represent information from similar subregions of the input space. The inhibition radius is dynamically adjusted to ensure local inhibition affects mini-columns with inputs from the same region of the input space. That is, *ϕ* increases if the average receptive field size increases. Specifically, *ϕ* is determined by the product of the average connected input spans of all SP mini-columns and the number of mini-columns per input. If SP inputs and mini-columns have the same dimensionality, *ϕ*=*γ* initially. In practice we also deal with input spaces that have no natural topology, such as categorical information (Purdy, 2016). In this case there is no natural ordering of inputs and we use an infinitely large *ϕ* to implement global inhibition.

Given an input pattern **z**, the activation of SP mini-columns is determined by first calculating the feedforward input to each mini-column, which we call the input overlap 
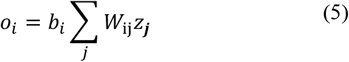

*b_i_* is a positive boost factor that controls the excitability of each SP mini-column.

A SP mini-column becomes active if the feedforward input is above a stimulus threshold *θ*_stim_and is among the top *s* percent of its neighborhood, 
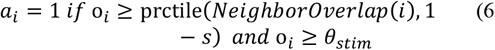

We typically set *θ*_stim_ to be a small positive number to prevent mini-columns without sufficient input to become active. prctile(∙, ∙) is the percentile function.*NeighborOverlap*(*i*) is the overlap values for all neighboring mini-columns of the *i*th mini-column. 
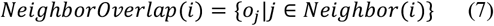

*s* is the target activation density (we typically use *s*=0.02). The activation rule (Eq. 6-7) implements *k-*winners-take-all computation within a local neighborhood. It has been previously shown that such computation can be realized by integrate-and-fire neuron models with precise spike timings (Billaudelle and Ahmad, 2015a). In this study we use discrete time steps to speed up simulation.

The feedforward connections are learned using a Hebbian rule. For each active SP mini-column, we reinforce active input connections by increasing the synaptic permanence by *p*^+^, and punish inactive connections by decreasing the synaptic permanence by *p*^-^. The synaptic permanences are clipped at the boundaries of 0 and 1.

To update the boost factors, we compare the recent activity of each mini-column to the recent activity of its neighbors. We calculate the time-averaged activation level for the each mini-column over the last *T* inputs as 
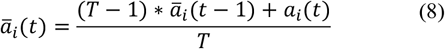

Where *a_i_(t)* is the current activity of the *i*th mini-column at time *t*. *T* controls how fast the boost factors are updated. Because the activity is sparse it requires many steps before we can get a meaningful estimate of the activation level. Typically we choose *T* to be 1000. The time-averaged activation level in Eq. 8 can be approximated by low-pass filtering of the voltage signal or intracellular calcium concentration. Similar calculations have been used in previous models of homeostatic synaptic plasticity (Clopath et al., 2010; Habenschuss et al., 2013).

The recent activity in the mini-column’s neighborhood is calculated as 
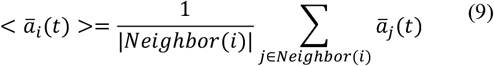

The boost factor *b_i_* is then updated based on the difference between *a_i_(t)* and < *ā_i_(t)*> as shown in Fig. 1D.
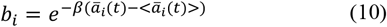

Here *β* is a positive parameter that controls the strength of the adaptation effect. The above boosting mechanism is inspired by numerous studies of homeostatic regulation of neuronal excitability (see (Davis, 2006) for a review). The mechanism encourages efficient use of mini-columns. The exact formula is not critical; we chose Eq. (10) due to its simplicity.

## 4. Properties of the HTM spatial pooler

In this section we describe a set of desirable properties for the HTM spatial pooler. These properties ensure flexible and robust representations of input streams with changing statistics, and are important for downstream neural computation.

The first property of the SP is to form fixed-sparsity representations of the input. To contribute to further neural computation, the outputs of the SP have to be recognized by downstream neurons. A cortical neuron recognizes presynaptic input patterns by initiating nonlinear dendritic spikes (Major et al., 2013) or somatic action potentials (Bean, 2007), with thresholds depending on intrinsic cellular properties. It has been previously shown that recognition of presynaptic activation patterns is robust and reliable if the presynaptic inputs have a fixed level of sparsity (Ahmad and Hawkins, 2016). However, if the sparsity is highly variable, input patterns with high activation densities would be more likely to cause dendritic spikes or action potentials in downstream neurons, whereas patterns with low activation densities would be much harder to detect. This will contribute to high false positive error for high density patterns and false negative error for low density patterns. A fixed sparsity is desirable because it ensures all input patterns can be equally detected.

A second desirable property is that the system should utilize all available resources to learn optimal representations of the inputs. From an information theoretic perspective, neurons that are almost always active and neurons that do not respond to any of the input patterns convey little information about the inputs. Given a limited number of neurons, it is preferable to ensure every neuron responds to a fraction of the inputs such that all neurons participate in representing the input space. The boosting mechanism in the SP (Eq. 8-10) is designed to achieve this goal. We quantify this property using an entropy metric (see details below).

A third desirable property is that output representations should be robust to noise in the inputs. Real-world problems often deal with noisy data sources where sensor noise, data transmission errors, and inherent device limitations frequently result in inaccurate or missing data. In the brain, the responses of sensory neurons to a given stimulus can vary significantly (Tolhurst et al., 1983; Faisal et al., 2008; Masquelier, 2013; Cui et al., 2016c). It is important for the SP to have good noise robustness, such that the output representation is relatively insensitive to small changes in the input.

A fourth property is that the system should be flexible and able to adapt to changing input statistics. The cortex is highly flexible and plastic. Regions of the cortex can learn to represent different inputs in reaction to changes in the input data. If the statistics of the input data changes, the spatial pooler should quickly adapt to the new data by adjusting its synaptic connections. This property is particularly important for applications with continuous data streams that has fast-changing statistics (Cui et al., 2016b).

Finally, a fifth property is that the system should be fault tolerant. If part of the cortex is damaged, as might occur in stroke or traumatic brain injury, there is often an initial deficit in perceptual abilities and motor functions which is followed by substantial recovery that occurs in the weeks to months following injury (Nudo, 2013). It has also been documented that the receptive fields of sensory neurons reorganize following restricted lesions of afferent inputs, such as retinal lesions (Gilbert and Wiesel, 1992; Baker et al., 2005). The spatial pooler should continue to function in the event of system faults such as loss of input or output neurons in the network.

## 5. Spatial pooler metrics

In addition to gauging performance in end-to-end HTM systems, we would like to quantify the performance of spatial pooler as a standalone component. Since the spatial pooler is an unsupervised algorithm designed to achieve multiple properties, we describe several statistical metrics that can be directly calculated based on the inputs and outputs of the spatial pooler. Such metrics are particularly useful if configurations of the spatial pooler caused the end-to-end HTM system to have poor performance.

### Metric 1: sparseness

We define the population sparseness as
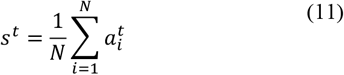
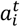 is the activity of the *i*th mini-column at time step t, *N* is the number of SP mini-columns. This metric reflects the percentage of active neurons at each time step. Since we consider binary activations (Eq. 6), the sparsity is straightforward to calculate. This metric has the same spirit as other population sparseness metrics for scalar value activations (Willmore and Tolhurst, 2001). We can quantify how well the spatial pooler achieves a fixed sparsity by looking at the standard deviation of the sparseness across time.

### Metric 2: entropy

Given a dataset of *M* inputs, the average activation frequency of each SP mini-column is
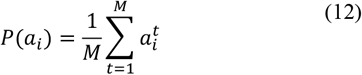

The entropy of the *i*th SP mini-column is given by the binary entropy function.
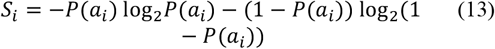

If *P*(*a_i_*) equals zero or one, we set *S_i_* to zero following convention. We define the entropy of the spatial pooler as 
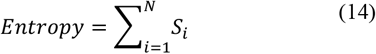

Since the average sparseness is almost constant in SP, the entropy is maximized when every SP mini-column has equal activation frequency. The spatial pooler will have low entropy if a small number of the SP mini-columns are active very frequently and the rest are inactive. Therefore, the entropy metric quantifies whether the SP efficiently utilizes all mini-columns.

### Metric 3: noise robustness

We test noise robustness by measuring the sensitivity of the SP representation to varying amount of input noise. We denote a clean input and a noise contaminated input as **z***_i_* and 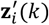 respectively, where *k*denotes the amount of noise added to the input. The corresponding SP outputs are denoted as **a***_i_* and 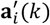 respectively. In our simulations, we randomly flip *k* percent of the active input bits to inactive, and flip the corresponding number of inactive input bits to active. This procedure randomizes inputs while maintaining constant input sparsity. We vary the amount of noise between 0 and 100%, and measure the fraction of shared active mini-columns in the SP output, averaged over a set of *M* inputs. The noise robustness index is defined as
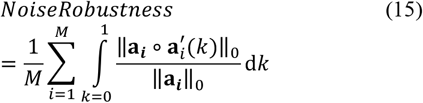

The L0-norm ||∙||_0_ gives the number of non-zero bits in a binary vector; the ∘ operator represents element-wise multiplication. Note that with binary vectors the L0-norm is identical to the L1-norm. The noise robustness index measures the area under the output overlap curve in Fig. 2C. The fraction of shared active mini-columns start at 100% when noise is zero, and decreases towards 0 as the amount of noise increases. The noise robustness thus lies between 0 and 1.

**Figure 2.**
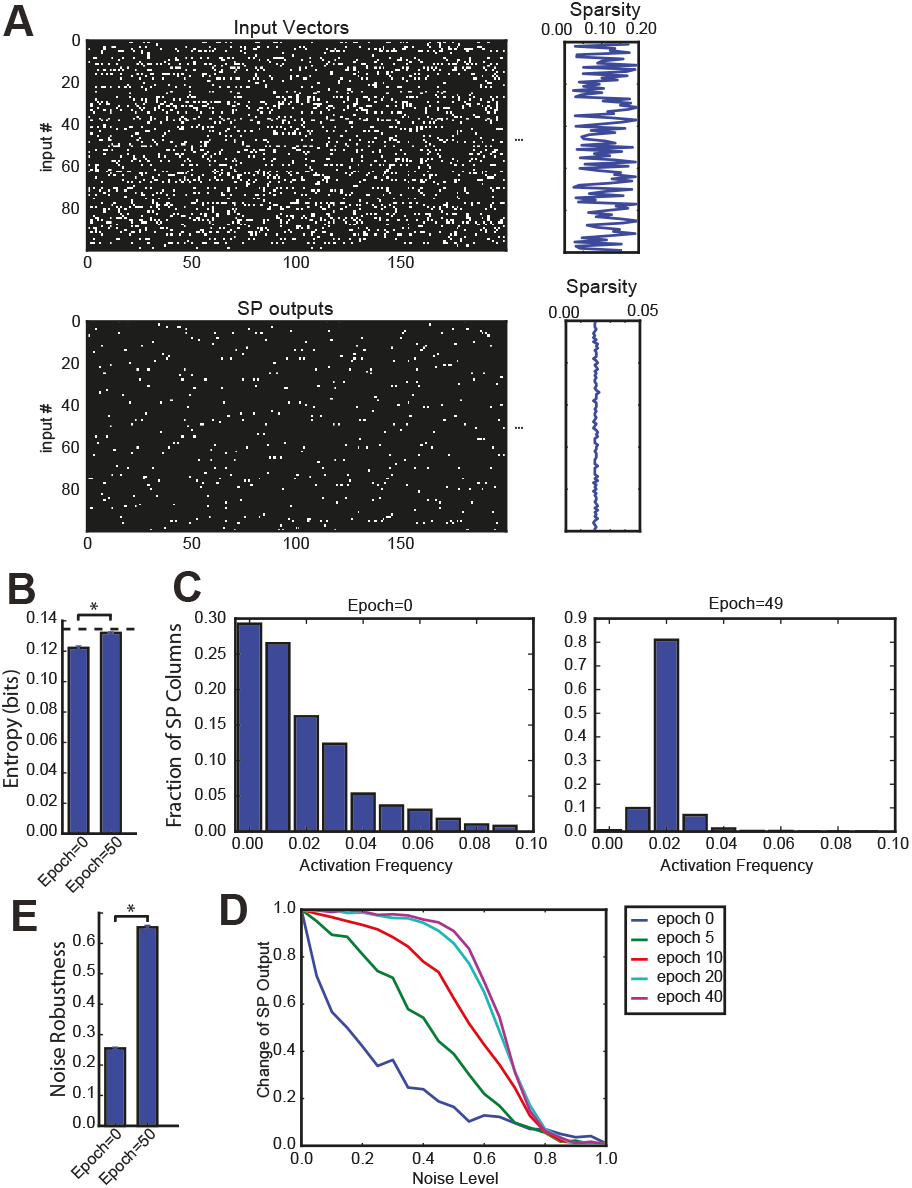
Spatial pooler forms SDRs with fixed sparsity and good noise robustness. **A.** We trained SP on a set of 100 randomly generated inputs (*top*). The input sparsity varies between 2% to 20%. The sparsity of the SP output lies close to 2% (*bottom*), despite the large variation of input sparsity. **B.**Entropy before and after learning, averaged across 10 repeated experiments (* *p*<10^-16^, *n*=10, paired t-test). The maximum possible entropy is shown as the black dashed line. **C.** The distribution of activation frequency of SP mini-columns. Before learning (*left*), a significant fraction of the SP mini-columns (~30%) are not being used at all, while other mini-columns are active much more frequently. After learning (*right*), almost every mini-column is active for 2% of the time, suggesting every SP mini-column participate in representing the input. As a result, the entropy of the distribution is much higher. **D-E** Noise robustness of SP. We tested SP on noisy inputs during learning. **D**. The change of the SP outputs is plotted as a function of the noise level. Before learning, a small amount of noise will lead to significant change in the SP output (*blue*), whereas after learning, there is almost no change in the SP output when 50% of the input bits changed. **E.** Average noise robustness before and after learning (* *p*<10^-16^, *n*=10, paired t-test).

### Metric 4: stability

Since the spatial pooler is a continuously learning system, it is possible that the representation for a given input changes over time or becomes unstable. Instability without changes in the input statistics could negatively impact downstream processes. We measure stability by periodically disabling learning and presenting a fixed random subset of the input data. The stability index is the average fraction of active mini-columns that remained constant for each input.

Denote the SP output to the *i*th test input at the *j*th test point as 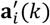, the stability index at test point *j* is given as,
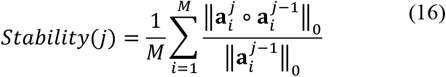

*M* is the number of inputs tested. The stability lies between 0 and 1, and equals 1 for a perfectly stable spatial pooler. Note that the superscript *j* is the index for test points instead of time steps in Eq. 16. We compute the percentage overlap between the SP outputs to the same test inputs across consecutive test points. We train the SP on the entire set of training data between test points.

## 6. Simulation details

We ran a number of different simulations (datasets described below). We used either a 2-dimensional spatial pooler with 32×32 mini-columns for experiments with topology, or a dimensionless spatial pooler with 1024 mini-columns for experiments without topology. The complete set of spatial pooler parameters is given in Table 1. The source code for all experiments are openly available at:https://github.com/numenta/nupic.research/tree/master/projects/sp_paper

**Table 1.** Parameters for the HTM spatial pooler.

| Common parameters | Value |
| --- | --- |
| Activation density | 2% |
| Connection threshold for synaptic permanence $\theta_c$ | 0.5 |
| Synaptic permanence increment $p^+$ | 0.1 |
| Synaptic permanence decrement $p^-$ | 0.02 |
| Boosting strength $\beta$ | 100 |
| Activation frequency duty cycle $T$ | 1000 |
| Stimulus threshold $\theta_{stim}$ | 1 |
| Fraction of potential inputs $p$ | 1 |
| Parameters for the 1-dimensional spatial pooler (no topology) |  |
| Column dimensions | 1024x1 |
| Potential input radius $\gamma$ | ∞ |
| Parameters for the 2-dimensional spatial pooler (with topology) |  |
| Column dimensions | 32x32 |
| Potential input radius $\gamma$ | 5 |

We presented each dataset in a streaming online fashion. Each dataset is presented to the SP in one or more epochs. We define an epoch as a single pass through the entire dataset in random order. Note that this definition of epoch is different from batch training paradigm because the SP receives one input pattern at a time and does not maintain any buffer of the entire dataset. We measured SP metrics between epochs on a random subset of the input data with learning turned off. In practice, the metrics could also be monitored continuously during learning.

We used the following datasets:

### Random Sparse Inputs

In this experiment we created a set of 100 random inputs with varying sparsity levels. Each input is a 32×32 image where a small fraction of the bits are active. The fraction of active inputs is uniformly chosen between 2% to 20%. This dataset employs a spatial pooler with topology and is used in Figs. 2-3.

**Figure 3.**
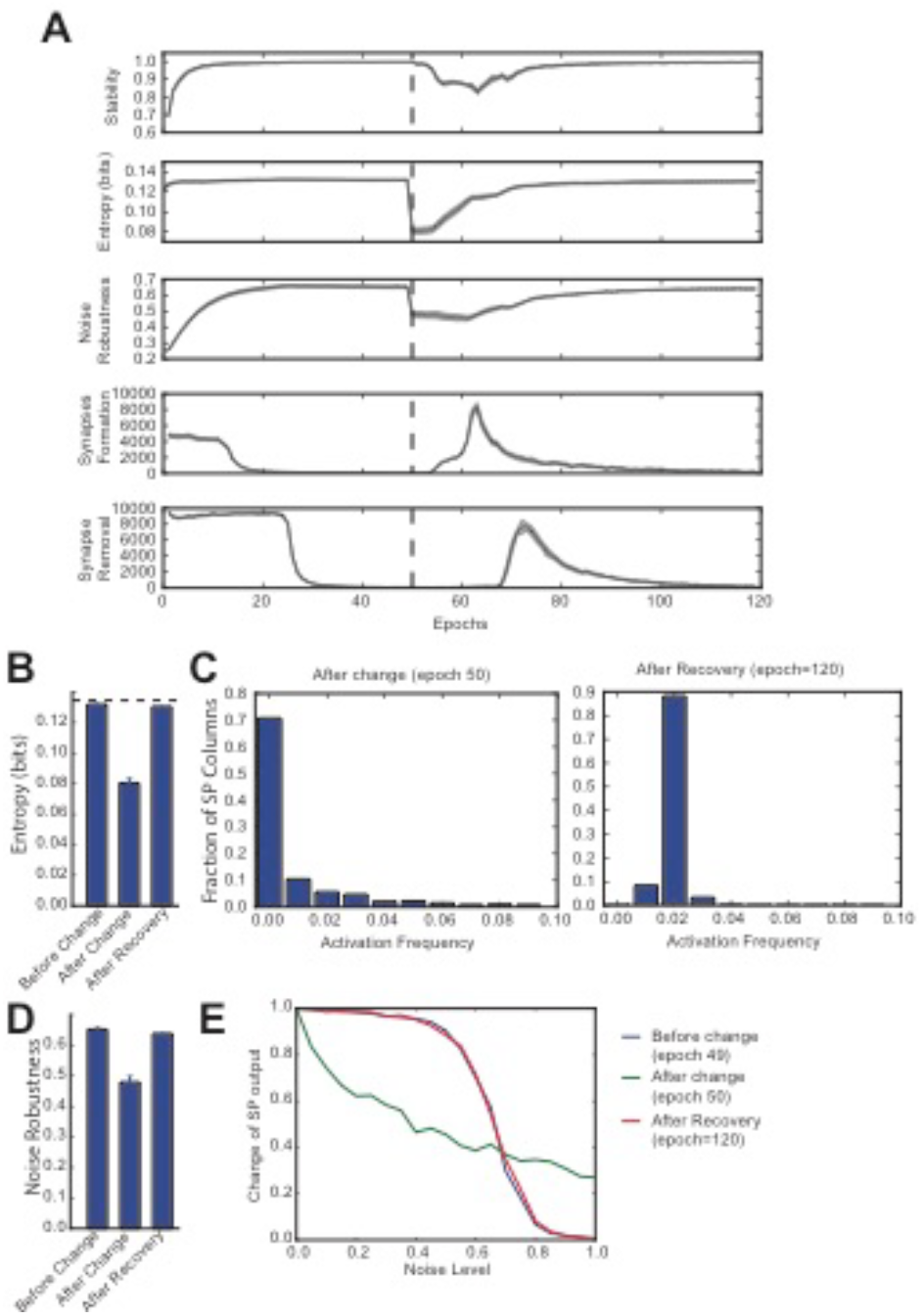
Continuous learning with HTM spatial pooler. SP continuously adapts to the statistics of the input data. SP is trained on a set of random inputs (described in Fig. 2) until it stabilizes. We then switch to a new set of inputs (black dashed line) and monitor the continuous adaptation of SP to the new dataset. **A.** Statistical metrics on SP during continuous learning: top: stability, 2nd row: entropy; 3rd row: noise robustness; 4th row: formation of new synapses; 5th row: removal of synapses. **B.** The entropy decreases right after the change of the input dataset (epoch=50), and recovers completely after the SP is trained on the new dataset for long enough time (epoch=120). The black dashed line showed the theoretical limit for entropy given the sparsity constraint. **C.** Distribution of activation frequency of SP mini-columns right after the change in dataset (left) and after recovery (right). **D**. Noise robustness before change (epoch=49), right after change (epoch=50) and after recovery (epoch=120). **E.** The noise robustness decreases right after the change of the input dataset (blue vs. green), and recovers completely after the SP is trained on the new dataset for long enough time (red).

### Random Bars and Crosses

The random bar dataset consists of 100 pairs of random bars. Each random bar pair stimuli is a 10x10 image, with a horizontal bar and a vertical bar placed at random locations. The bars have a length of 5 pixels. Each random cross stimuli is a 10x10 image with a single cross. The random cross dataset consists of 100 cross patterns at random locations, where each cross consists of a horizontal bar and a vertical bar that intersect at the center. Each bar has a length of 5 pixels. This dataset is used in Fig. 4A-B.

**Figure 4.**
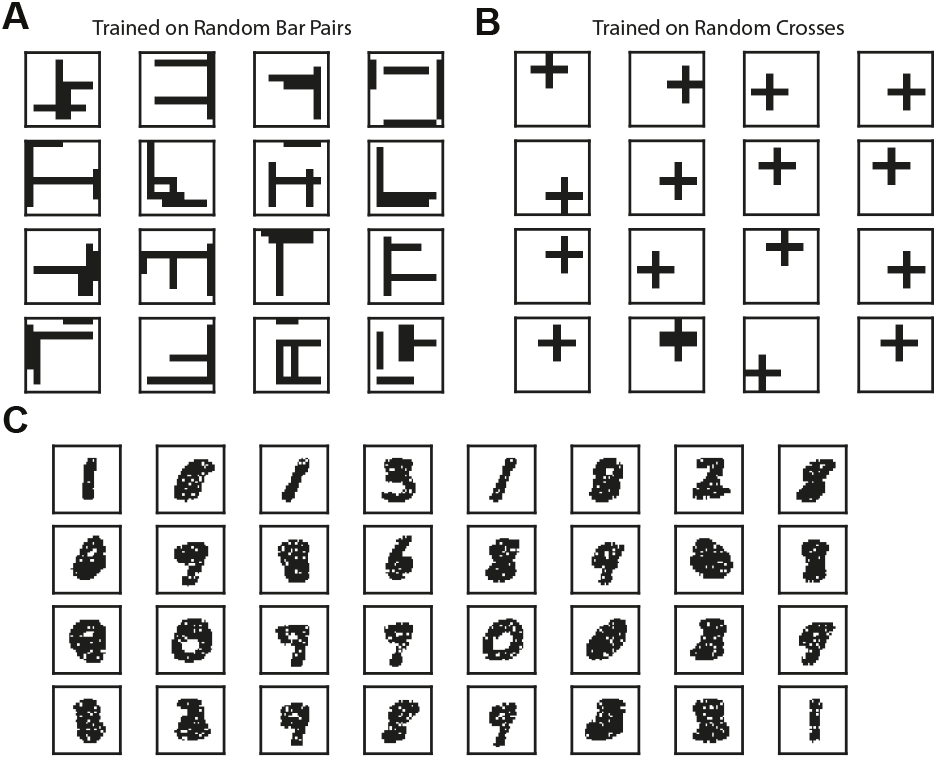
Example receptive fields of SP. The receptive fields of SP mini-columns capture statistics of the input data. We define receptive field as the set of inputs that are connected to a mini-column. **A.** Example SP Receptive fields trained on random bar pairs. **B.** Example SP receptive fields trained on random crosses. **C.** Example SP receptive fields trained on MNIST dataset.

### MNIST

We trained a spatial pooler without topology on the MNIST database of handwritten digits (Lecun et al., 1998). We present the full training set of 60,000 examples in a single epoch to the spatial pooler. We visually examined the receptive field structures of a subset of randomly selected SP mini-columns after training. This dataset is used in Fig. 4C.

### Fault tolerance with topology

For the fault tolerance experiment (Fig. 5), we used images of random bar sets as input. The input space has dimensionality of 32×32 and each input contains 6 randomly located horizontal or vertical bars. Each bar has a length of 7. We used an SP with a two-dimensional topology and 32×32 mini-columns. We first trained the intact SP on the random bars input until it stabilized (after 18000 inputs). We then tested two different types of trauma: simulated stroke or simulated input lesion. During the simulated stroke experiment, we permanently eliminated 121 SP mini-columns that lie in an 11x11 region at the center of the receptive field. For the simulated input lesion experiment, we did not change the SP during the trauma. Instead, we permanently blocked the center portion of the input space (121 inputs that lie in an 11x11 region at the center of the input space). For both experiments, we monitored the recovery of the SP for another 42000 steps.

**Figure 5.**
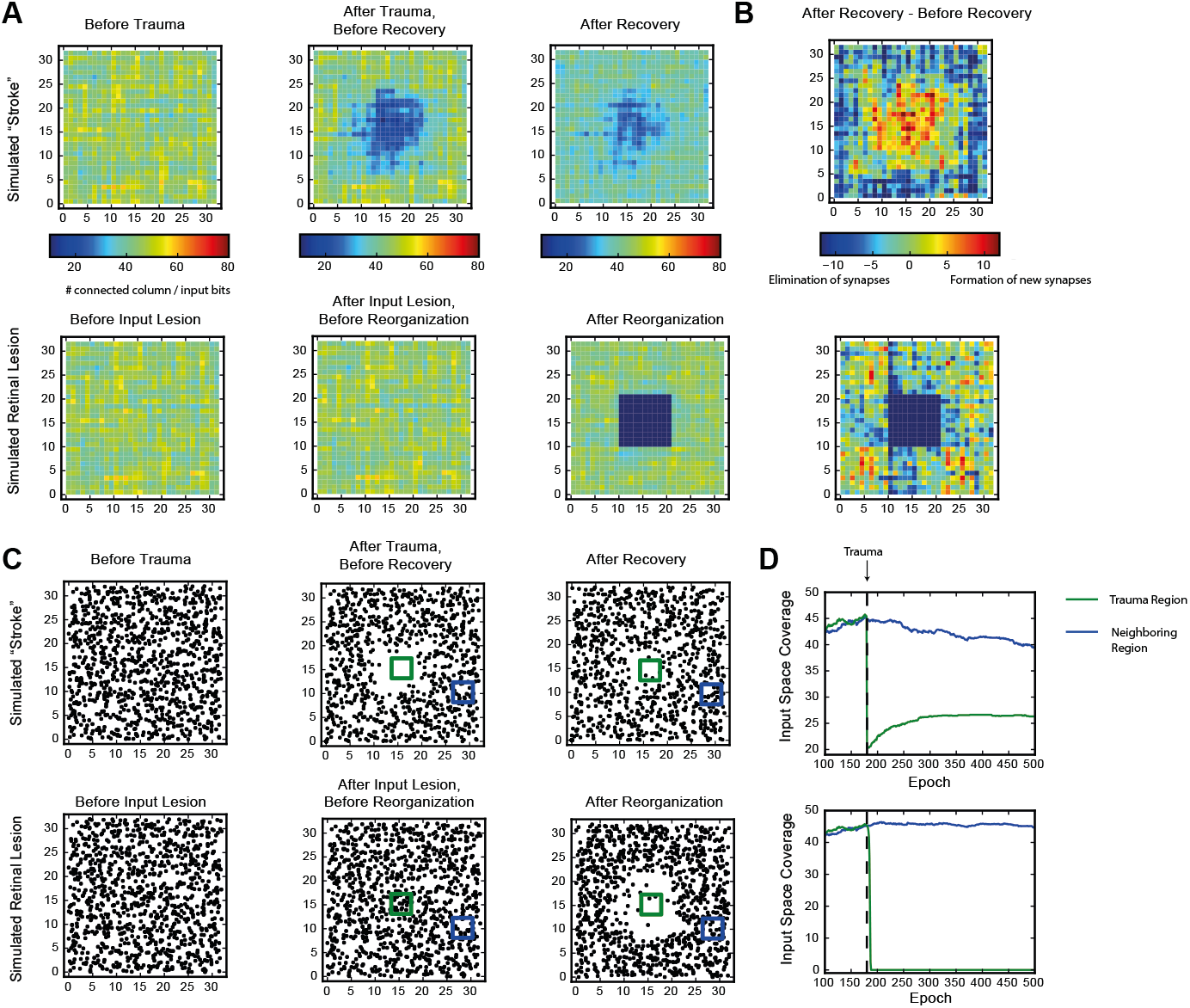
Recovery of HTM spatial pooler after damage and input lesion. During the simulated stroke, a fraction of SP mini-columns that are connected to the center region of the input space is killed. During the simulated retinal lesion, the center portion of the input space is blocked while the spatial pooler and its feedforward inputs are kept intact. **A.** The number of SP mini-columns connected to each input bits before trauma (*left*), right after trauma (*middle*) and after recovery (*right*). The simulated stroke experiment is shown at the top and the simulated retinal lesion experiment is shown at the bottom. **B.** Growth and elimination of synapses during the recovery process for the simulated stroke (*top*) and retinal lesion (*bottom*) experiment. **C.** Receptive field centers of all SP mini-columns before trauma (*left*), right after trauma (*middle*) and after recovery (*right*). **D.** Number of mini-columns connected to the center region (green square in Fig. 5C) and a neighboring region (blue square) during the recovery process. The recovery is very fast for the retinal lesion experiment (bottom), and slower for the simulated stroke experiment (top).

### NYC taxi passenger count prediction

In addition to the above artificial datasets, we also tested the SP in an end-to-end real-world HTM system. We chose the problem of demand prediction for New York City taxis. The dataset is publicly available via the New York City Metropolitan Authority. Full details are described in our previous paper (Cui et al., 2016a). As described in Fig. 1, the input data stream is first converted to binary representations using a set of encoders (Purdy, 2016). The spatial pooler takes the output of the encoders as input and forms sparse distributed representations. The HTM sequence memory then learns sequences of SDRs and represents sequences with a sparse temporal code. Finally, we use a single layer feedforward classification network to map outputs of HTM sequence memory into real-time predictions for future inputs. The task is to model a continuous stream within the context of a real-time application. As such the spatial pooler, temporal memory, and classifier all learn continuously. It is important for the spatial pooler to output robust and efficient representations in order for the downstream components to learn.

Following (Cui et al., 2016a), we aggregated the passenger counts in New York City taxi rides at 30-minute intervals. We encoded the current passenger count, time of day and day of week into binary vectors using scalar and date-time encoders (Purdy, 2016). A spatial pooler with global inhibition was trained continuously on the outputs of encoders and provided input to the HTM sequence memory (Cui et al., 2016a). To evaluate the role of learning and boosting in SP, we compared the prediction accuracy in three scenarios (1) SP without learning or boosting, (2) SP with learning but not boosting, and (3) SP with both learning and boosting.

## 7. Results

We first discuss results on the Random Sparse Inputs dataset with respect to the metrics (Figs. 2 and 3). The input patterns are presented repeatedly to the HTM spatial pooler in a streaming fashion. In our simulation, the population sparsity of the spatial pooler is always close to the target level of 2%, even though the input sparsity varies widely in the range of 2%~20% (Fig. 2A). This is an inherent property of the network due to the use of local *k*-winners-take-all activation rules.

We measured the average entropy across all mini-columns (see Spatial Pooler Metrics). Since the overall activation sparsity is fixed in our network, the entropy is maximized if all mini-columns have the same activation probability. In this experiment, the entropy increases from 0.1221±0.0013 bits/mini-column to 0.1320±0.0007 bits/mini-column with training. The difference is highly significant across repeated experiments with different set of random inputs (*p*<10^-8^, *n*=10, paired t-test). As a reference, the maximum possible entropy is 0.1345 bits/mini-column with the same sparsity levels. The increase of entropy is due to efficient use of all mini-columns. Before learning, a significant fraction of the mini-columns (~30%) were not active for any of the input, whereas a small fraction of the mini-columns were much more active than others. After learning, almost every mini-column was active for 2% of the time.

Figs. 2D and 2E demonstrate noise robustness as a function of SP learning. Before learning, a small change in the input will cause a large change in the SP output, suggesting high noise sensitivity (Fig. 2D, blue). After learning, the noise robustness gradually improves. After 40 repetitions of the dataset, there is no change on the SP output even if 40% of the active input bits are changed. The average noise robustness index (Eq. 15) correspondingly improves from 0.254±0.004 to 0.652±0.007 (Fig. 2E, *p*<10^-16^, *n*=10, paired t-test). The improved noise robustness is due to the Hebbian learning rules. A set of SP mini-columns forms reliable connections to active input neurons during learning. The same set of SP mini-columns can be activated even if some of the input neurons are affected by noise after learning.

To test whether the spatial pooler can adapt to changing inputs, we train the spatial pooler until it stabilizes on one set of inputs. We then present a completely different input dataset (Fig. 3A). Right after we switch to a new dataset, the entropy and noise robustness drops sharply (Fig. 3B, 3D). At this point a large fraction of the SP mini-columns are not responsive to any input in the new dataset (Fig. 3C, left). Once learning resumes the spatial pooler quickly adapts to the new input dataset and the performance metrics recover back to the levels before the change (Fig. 3B, 3D). The SP adapts to the new dataset by first forming many more new synapses (Fig. 3A, 4th row) and then pruning unnecessary connections later (Fig. 3A, 5th row).

The spatial pooler achieves these properties by continuously adapting feedforward connections to the input data. To illustrate how receptive field structures of SP are shaped by the input data, we trained the spatial pooler on the Random Bars dataset. In this experiment, the training data consists of either pairs of randomly generated bars, or a single cross at a random location.

We plot example receptive fields from a random subset of SP mini-columns in Fig. 4. For this dataset the receptive field typically contains horizontal and vertical bar structures at random locations (Fig. 4A). Each SP mini-column responds to more than one bar in order to achieve the target activation frequency of 2%. When trained on the random cross data stream, the resulting receptive field consists of cross structures that resemble statistics of the inputs (Fig. 4B). Note that the resulting receptive field also depends on the ratio of synaptic permanence decrement and increment (*p*^-^/*p*^+^). If we increase *p*^-^ to 0.05, most of the receptive field contains single bar segment when trained on the random bar pairs dataset (data not shown).

We also trained the spatial pooler on the MNIST dataset. In this case the receptive field contains digit-like structures (Fig. 4C). Although some receptive fields clearly detect single digits, others are responsive to multiple digits. This is because individual SP mini-columns do not behave like “grandmother cells”–they are not meant to detect single instances of the inputs. Instead a single input is collectively encoded by a set of SP mini-columns. When trained on complex natural datasets, we expect to see a diversity of receptive structures within the spatial pooler, which may not resemble specific input instances.

### 7.1. Fault tolerance

We evaluated whether the HTM spatial pooler has the ability to recover from lesion of the afferent inputs (input lesion experiment) or damage to a subset of the SP mini-columns (stroke experiment).

In the stroke experiment, the center portion of the input space becomes much less represented right after the trauma because the corresponding SP mini-columns are eliminated. The network partially recovers from the trauma after a few hundred epochs, with each epoch consisting of 100 inputs. During the recovery process, SP mini-columns near the trauma region shift their receptive field toward the trauma region and start to represent stimuli near the center (Fig. 5C, supplemental movie 1). The network forms many more new synapses in the center, which is accompanied by loss of synapses in the non-trauma region (Fig. 5B, top). There is a clear recovery on the coverage of the trauma region (Fig. 5D, bottom).

In the input lesion experiment, the center portion of the input space is blocked. Since there is no change on the SP mini-columns or the associated synaptic connections, there is no immediate change on the input space coverage (Fig. 5A, bottom) or the receptive field center distributions (Fig. 5C, bottom). However, the SP mini-columns quickly reorganize their receptive field within a few epochs. The SP mini-columns that respond to the center inputs starts to respond to inputs on the surrounding, non-damaged areas (Fig. 5C, supplemental movie 2). Almost all the connections to the lesion region are lost after the reorganization (Fig. 5B, bottom).

These demonstrate the fault tolerance and flexibility of the SP. The fixed sparsity and the homeostasis excitability control mechanism of the SP ensure that the input space is efficiently represented by all SP (undamaged) mini-columns. It is interesting to note that the different recovery speeds from the two simulations coincide with experimental studies. It has been reported that after focal binocular retinal lesions, the receptive field sizes increases within a few minutes for cortical neurons that lie near the edge of the retinal scotoma (Gilbert and Wiesel, 1992). In contrast, if part of the cortex is damaged, the recovery is partial and occurs on a much slower time scale (Nudo, 2013).

### 7.2. The spatial pooler in a real-world streaming analytics task

In this section we evaluate the role of the SP in an end-to-end real-world HTM system. We consider the problem of real-time prediction of the number of taxi passenger in New York City (Fig. 6A, see Methods). We have previously shown that HTM systems with a fixed pre-trained spatial pooler achieves state-of-the-art performance on this task (Cui et al., 2016b). Here we consider the role of learning in the spatial pooler and evaluate three scenarios: using a randomly initialized SP without learning, allowing SP learning but without boosting, and using a SP with both continuous learning and boosting.

**Figure 6.**
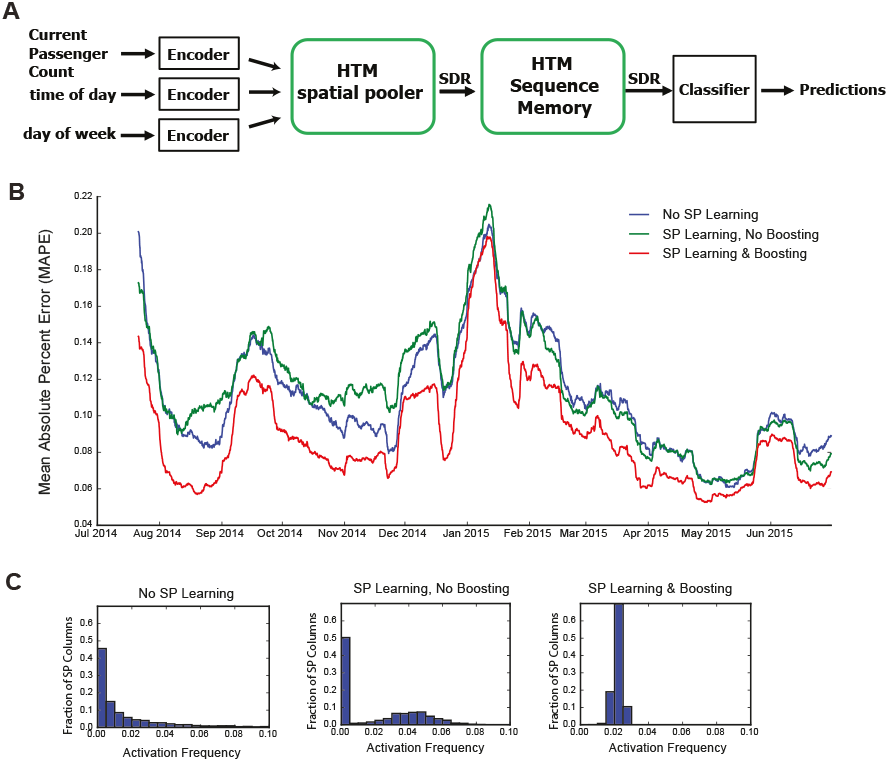
The role of HTM spatial pooler in prediction task. A. A complete HTM system of encoder -> spatial pooler -> sequence memory -> classifier is used for predicting the NYC taxi passenger count. B. Prediction error with an untrained random SP (blue), a SP with continuous learning but without boosting (green), and a SP with both continuous learning and boosting (red). C. Distribution of activation frequency of SP mini-columns. A large fraction of SP mini-columns are not being used for an untrained SP or for a SP without boosting. In contrast, almost all mini-columns are active for about 2% of the time when boosting is enabled.

We fed the inputs in a single pass to mimic the scenario of real-time online prediction. The time-averaged prediction error for the three cases is plotted as a function of training time (Fig. 6B). At the beginning of learning, the prediction error rapidly decreases, representing the initial learning phase of the system. The occasional increases in error reflect real world changes that correspond to events and holidays (e.g., Thanksgiving, Christmas, New York City Marathon, etc.).

The spatial pooler with both learning and boosting achieves the best performance throughout the prediction task. The spatial pooler with learning but without boosting is roughly comparable to a random static SP. This suggests the importance of both continuous Hebbian learning and homeostatic excitability control. The difference in performance can be understood by observing the distribution of activation frequency across SP mini-columns. Without the homeostatic excitability control, a large fraction of the SP mini-columns are not being used at all (Fig. 6C). It is more error-prone for the HTM sequence memory to learn transitions of such ill-behaved SDRs.

## 8. Discussion and Conclusions

In this paper, we described properties of the HTM spatial pooler, a neurally inspired algorithm for learning sparse distributed representations online. Inspired by computational principles of the neocortex, the goal of the HTM spatial pooler is to create SDRs and support essential neural computations such as sequence learning and memory. The model satisfies a set of important properties, including tight control of output sparsity, efficient use of mini-columns, preserving similarity among inputs, noise robustness, fault tolerance and fast adaptation to changes. These properties are achieved using competitive Hebbian learning rules and homeostatic excitability control mechanisms. We demonstrate the effectiveness of SP in an end-to-end HTM system on the task of streaming data prediction. The HTM spatial pooler leads to a flexible sparse coding scheme that can be used in practical machine learning applications.

### 8.1. Relationship with other sparse coding techniques

The spatial pooler learns sparse distributed representations for inputs. It is related to the broad class of sparse coding techniques, which uses activation of a small set of neurons to encode each item. One theory of sparse coding suggests that sparse activations in sensory cortices reduce energy consumption of the brain while preserving most of the information (Földiák, 2002; Olshausen and Field, 2004). Early studies of sparse coding explicitly optimize a cost function that combines low reconstruction error and high sparseness (Olshausen and Field, 1996a, 1997). When applied to natural images, these techniques lead to receptive fields that resemble those of V1 neurons (Olshausen and Field, 1996a; Lee et al., 2006), suggesting that the functionality of early sensory neurons can be explained by the sparse coding framework. Sparse coding has been implemented previously with biologically plausible local learning rules. Földiák showed that a neural network could learn a sparse code using Hebbian forward connections combined with a local threshold control mechanism (Földiák, 1990). It has been recently shown that such learning rules can be derived analytically (Hu et al., 2014).

Many of the properties we analyzed in this paper have also been discussed in previous studies of sparse coding. It has been shown that sparse representations are naturally robust to noise and can be used for robust speech recognition (Sivaram et al., 2010; Gemmeke et al., 2011), robust face recognition (Wright et al., 2009) and super resolution image reconstruction (Jianchao Yang et al., 2010). Online sparse coding and dictionary learning techniques have been proposed in previous studies in order to handle dynamic datasets (Mairal et al., 2010). It is known that representations learned from traditional sparse coding techniques have low entropy, as the probability distribution of activity of an output unit is peaked around zero with heavy tails (Olshausen and Field, 1996b). In this study we found that although the entropy is low compared to dense representations, it increases with training in the HTM spatial pooler. This is because the homeostatic excitability control mechanism encourages neurons in the SP to have similar activation frequencies, thus increasing the representational power of the network.

Most previous studies propose the goal of sparse coding is to avoid information loss, reduce energy consumption, and form associative memory with minimum cross talk (Olshausen and Field, 2004). A commonly used criterion is how well one can reconstruct the inputs given the sparse activations and a set of learned basis vectors. Although these studies explain receptive structure in primary visual cortex and lead to practical machine learning algorithms for feature selection (Gui et al., 2016) and data compression (Pati et al., 2015), the purpose of neural computation is more than preserving information. In this paper, we take a different perspective and ask how computational properties of the HTM spatial pooler contribute to downstream cortical processing in the context of HTM systems. Instead of reconstruction error, we define the performance of SP in terms of a set of properties, including population entropy, noise robustness, stability, and fault tolerance. It is important to perform a multi-dimensional assessment of the SP in order to ensure that it forms robust sparse distributed representations that capture semantic similarity of the inputs. We demonstrated that the HTM spatial pooler achieves these properties and that these properties contribute to improved performance in an end-to-end system. It is important to note that no single metric is sufficient to ensure SP is behaving properly. For example, one can achieve good noise robustness by always using a small set of SP mini-columns, but that will give bad entropy. It is easy to achieve high entropy by using a random output at each time step, but that will cause bad stability. It is important to consider all the metrics together. As a result, the learning algorithm of SP cannot be easily derived by optimizing a single objective function.

The Hebbian learning rules of HTM spatial pooler resemble many previous sparse coding algorithms (Földiák, 1990; Zylberberg et al., 2011; Hu et al., 2014) and associative memory models (Willshaw et al., 1969; Hecht-Nielsen, 1990; Bibbig et al., 1995). There are several differences. First, we include homeostatic excitability control as a gain modulation mechanism. The role of homeostasis is to make sure that the distribution of neural activity is homogeneous. It has been previously proposed that homeostasis is crucial in providing an efficient solution when learning sparse representations (Perrinet, 2010). Some models of synaptic plasticity do include homeostatic components in the learning rule that control the amount of synaptic weight change (Clopath et al., 2010; Habenschuss et al., 2013). The homeostatic excitability regulation mechanism in the SP achieves a similar effect without directly affecting the synapse modification process. Second, we use a local inhibition circuit that implements *k*-winners-take-all computation to have tight control over the output sparseness. This is important when SP activations are used by downstream neurons with dendrites that have threshold nonlinearities. Finally, we use binary synapses and learning via synaptogenesis (Zito and Svoboda, 2002). The use of binary synapses can dramatically speed up the computation. Overall the HTM spatial pooler algorithm is a suitable candidate for learning sparse representations online from streaming data.

### 8.2. Potential Neural Mechanisms of spatial pooler

A layer in a HTM system contains a set of mini-columns. Each mini-column contains cells with the same feedforward receptive field. The mini-column hypothesis has been proposed for several decades (Mountcastle, 1997; Buxhoeveden, 2002), but the utility of mini-columns remains controversial due to a lack of theoretical benefit (Horton and Adams, 2005). According to HTM theory cells within the same mini-column have the same feedforward connections but different lateral connections, thus representing the same feedforward input in different temporal contexts (Hawkins and Ahmad, 2016). We propose the spatial pooler models feedforward receptive field learning at the mini-column level. Experimental studies have shown that neurons within the same mini-column have almost identical receptive field locations, sizes and shapes, whereas RFs of neurons in neighboring mini-columns can differ significantly (Jones, 2000). This variability cannot be explained by the difference in feedforward inputs, because the extent of arborization of single thalamic afferent fibers, which can be as much as 900 µm in cats (Jones, 2000), is significantly more extensive than the dimensions of minicolumns, which typically have diameters around 20-60 µm (Buxhoeveden, 2002). (Favorov and Kelly, 1996; Jones, 2000). It requires dedicated circuitry mechanism to ensure that cells in the same mini-column acquire the same receptive field.

We propose two possible neural circuit mechanisms for the spatial pooler and discuss their anatomical support. In the first proposal (Fig. 7B, left), the feedforward thalamic inputs innervate both excitatory pyramidal neurons as well as an inhibitory neuron (*green*). This inhibitory neuron can indirectly activate the pyramidal neurons through a disynaptic dis-inhibition circuit. It acts as a “teacher” cell that guides the receptive field formation of excitatory neurons. There are many distinct classes of inhibitory neurons in the cortex. Some classes, such as bipolar cells and double bouquet cells exclusively innervate cells within a cortical mini-column (Markram et al., 2004; Wonders and Anderson, 2006). It is well documented that feedforward thalamacortical input strongly activates specific subtypes of inhibitory neurons (Gibson et al., 1999; Porter et al., 2001; Swadlow, 2002; Kremkow et al., 2016). It is possible these inhibitory neurons participate in defining and maintaining the feedforward receptive field of cortical mini-columns.

**Figure 7.**
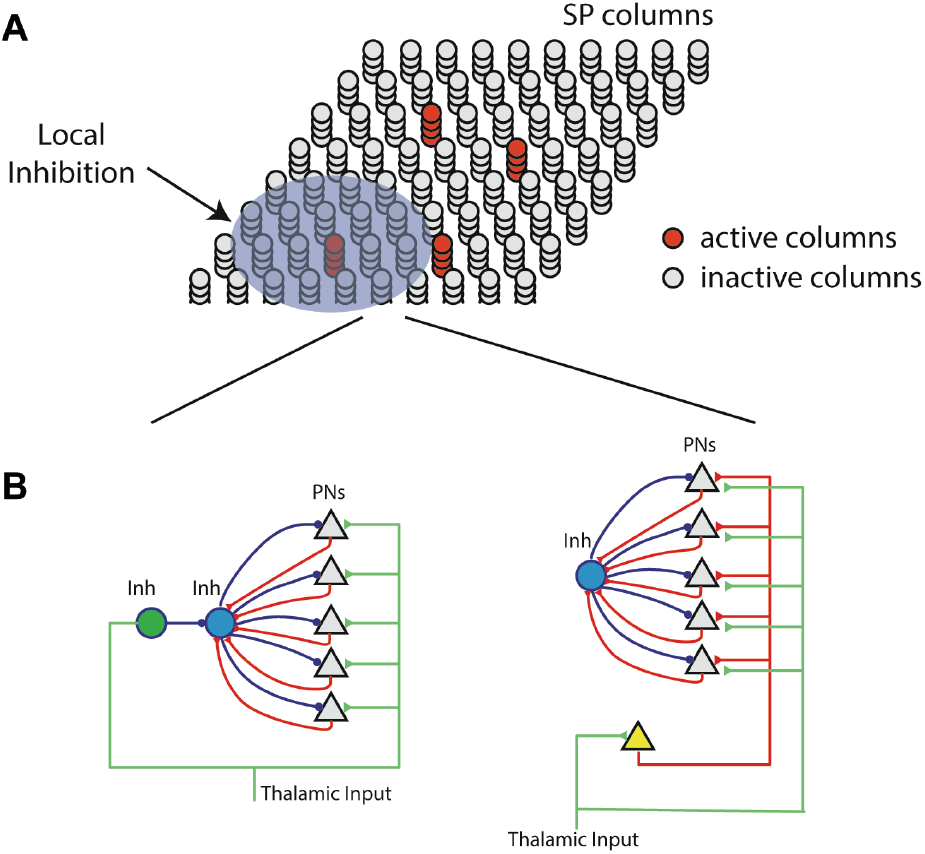
Neural mechanism of HTM Spatial pooler. **A.** Spatial pooler requires local across mini-column inhibition to ensure that a small fraction of the mini-columns are active at any time. **B.** Potential mechanisms to ensure neurons within the same mini-column share the same feedforward receptive field. *left*: the green inhibitory neuron controls the receptive fields of excitatory pyramidal neurons (gray triangles) through a dis-inhibition circuit. *right*: A single (or small number of) excitatory neurons (yellow) controls the receptive field of excitatory neurons. In both cases, PNs indirectly inhibit other PNs in the same mini-column through the within mini-column inhibition (blue).

In the second proposal, a single excitatory cell receives thalamic inputs and innervates all excitatory cells in a mini-column. This excitatory neuron guides the receptive field formation of other excitatory cells in the mini-column. A similar circuit has been observed during early development. Subplate neurons, a transient population of neurons, receive synaptic inputs from thalamic axons, establishing a temporary link between thalamic axons and their final targets in layer IV (Friauf et al., 1990; Ghosh and Shatz, 1992; Kanold et al., 2003). It remains to be tested whether a similar circuit exists in adult brain.

The spatial pooler relies on several other neural mechanisms. The learning rule is based on competitive Hebbian learning. Such learning can be achieved in the brain via synaptic plasticity rules such as long-term potentiation (Teyler and DiScenna, 1987), long-term depression (Ito, 1989), or spike-time dependent plasticity (Song et al., 2000). Homeostatic excitability control mechanisms, analogous to the spatial pooler’s boosting rule, have been observed in cortical neurons (Davis, 2006). Finally, the *k*-winners-take-all computation in the SP can be implemented using leaky integrate-and-fire neuron models (Billaudelle and Ahmad, 2015b).

## 9. Acknowledgement

We thank the editor and the reviewers for their constructive comments and suggestions. We thank Mirko Klukas for suggestions on the entropy metric. We also thank the NuPIC open source community (numenta.org) for continuous support and enthusiasm about HTM.

